# Crystal structure and molecular dynamics simulations of rademikibart Fab-IL-4Rα complex reveal biochemical basis for next-generation potent IL-4Rα inhibition in type 2 allergic and inflammatory diseases

**DOI:** 10.64898/2026.04.12.718052

**Authors:** Yuanjun Shi, Kelsey Nolden, Minh Ho, Haote Li, Victor S. Batista, Raúl Collazo, Christopher G. Bunick

## Abstract

Rademikibart (CBP-201) is a human monoclonal antibody with higher binding affinity to IL-4Rα compared to dupilumab. Dupilumab is a first-generation interleukin-4 receptor alpha (IL-4Rα) inhibitor for treating IL-4Rα-dependent inflammatory disorders, including several dermatologic and respiratory conditions. Rademikibart, however, demonstrated better inhibition of STAT6 intracellular signaling *in vitro* and similar potency in inhibiting both IL-4 induced TARC release and IL-4 induced B cell activation. To further characterize the molecular function of rademikibart and its differentiation from dupilumab, we determined the crystal structure of the rademikibart fragment antigen binding (Fab) bound to IL-4Rα at 2.71 Å resolution and compared this to the 2.82 Å resolution structure of dupilumab Fab bound to IL-4Rα. The rotation angle between dupilumab and rademikibart bound to IL-4Rα is 54.88°. This rotation enables the binding epitopes of rademikibart, but not dupilumab, on IL-4Rα to overlap more closely with the conserved binding interface naturally utilized by IL-4 and IL-13 cytokines. Molecular dynamics (MD) studies on rademikibart and dupilumab bound to IL-4Rα examined the stability of the complexes and effects of amino acid mutations on receptor complex formation. MD simulations demonstrated that the third interface loop (residues 145 to 153 in domain 2) of IL-4Rα interacts directly with rademikibart, which is absent in the dupilumab/IL-4Rα complex. This finding is confirmed by increased hydrogen bond interactions at the interface between rademikibart and IL-4Rα, demonstrating superior binding energy for rademikibart. Through analysis of the x-ray crystallography structures, MD-equilibrated structures, and computational point-mutation analysis of rademikibart, we identified residue Y50 and R55 of the light chain and R97, R99, and Y101 of the heavy chain of rademikibart as key residues interacting with IL-4Rα’s third interface loop. Our data provides a molecular and structural rationale for the enhanced IL-4Rα inhibition by rademikibart over dupilumab, confirming rademikibart as an optimized second-generation IL-4Rα inhibitor.

## Introduction

Targeted biologics have reshaped the therapeutic paradigm for type 2 (T2) inflammatory diseases, shifting care from broad immunosuppression to pathway-specific interventions. Biologics targeting IL-4 signaling have become critical in managing these T2 immune-mediated disorders, as IL-4 is a key driver of the pathway, initiating Th2 cell differentiation, promoting IgE class switching, and amplifying cytokine signaling that sustains allergic and inflammatory responses across multiple tissues.

The first global approval of an anti-IL-4 biologic, dupilumab, occurred in 2017 when the U.S. FDA authorized dupilumab for the treatment of adults with moderate-to-severe atopic dermatitis (AD), marking the first targeted biologic for AD and inaugurating a new era of mechanism-based care. Subsequent indications extended dupilumab’s reach across T2 inflammatory diseases including, but not limited to, asthma^1^, chronic rhinosinusitis with nasal polyposis^2^, and chronic obstructive pulmonary disease^3–5^. These approvals have materially altered standards of care by reducing reliance on systemic corticosteroids, elevating expectations for improved efficacy, providing sustained disease control and improved patient-reported outcomes across both dermatology and respiratory medicine, and improving long-term safety. However, dupilumab’s use is associated with notable adverse events, including injection-site reactions, conjunctivitis, facial erythema, eosinophilia, and rare systemic complications^6–15^. These effects, though generally manageable, highlight the complexity of modulating IL-4/IL-13 signaling and the need for therapies that maintain efficacy longer while improving tolerability.

Rademikibart is a next-generation IL-4Rα inhibitor that aims to address these challenges through optimized molecular design and is currently undergoing phase 2 trials for the treatment of both acute asthma and COPD (NCT06940141 and NCT06940154). Rademikibart and dupilumab are both fully human IgG4κ monoclonal antibodies that bind IL-4Rα. Rademikibart demonstrated binding to a distinct IL-4Rα epitope with higher affinity than dupilumab in in-vitro and ex-vivo studies, indicating a similar mechanism with potential biophysical differentiation at the receptor interface^16^. By refining epitope engagement and reducing variability in immune modulation, rademikibart may offer a differentiated safety profile and improved patient experience, reinforcing its role as an evolution in IL-4Rα-targeted therapy.

At the disease-system level, dual IL-4/IL-13 inhibition broadly suppresses T2 inflammation, consistent with clinical benefits observed across various dermatological and respiratory diseases. For instance, while IL-13 contributes prominently to epithelial remodeling (e.g., mucus biology) in the airways and barrier dysfunction in skin, IL-4 is pivotal for initiating and sustaining T2 immunity—driving Th2 polarization, class-switching to IgE, and multiorgan amplification of allergic inflammation^17,18^. Consequently, blocking IL-4Rα (rather than IL-13 alone) targets the common upstream node governing both cytokines’ signaling, producing broader disease-modifying potential across dermatologic and respiratory manifestations^19,20^.

From a structural standpoint, IL-4 and IL-13 are four α-helix bundle cytokines with a small, antiparallel β-sheet, which signal through overlapping receptor complexes^21–23^. IL-4 uniquely engages both the type I receptor (IL-4Rα/γc) and the type II receptor (IL-4Rα/IL-13Rα1), whereas IL-13 signals primarily through the type II receptor. Although the type II receptor is structurally identical in both cases, differences in sequence and thermodynamic properties influence how the heterotrimeric receptor–ligand complex assembles, depending on whether IL-4 or IL-13 is binding^22^. This receptor architecture makes anti-IL-4Rα blockade foundational, because targeting IL-4Rα simultaneously prevents IL-4 and IL-13–dependent assembly and downstream JAK–STAT (STAT6) signaling; therefore, IL-4Rα serves as an upstream choke point for T2 immune amplification relevant to dermal and airway tissues^19^.

Understanding this structural and thermodynamic complexity sets the stage for examining how IL-4Rα mediates cytokine binding at the receptor interface, which is a critical determinant of signaling potency and a key target for precise therapeutic intervention. The IL-4Rα subunit of type I and type II IL-4/IL-13 receptors engages either IL-4 or IL-13 through a conserved mechanism and shared epitope (**Figure 1**). The IL-4Rα subunit binds IL-4/IL-13 via three major loops belonging to two separate protein domains (D1: loops 2 and 3; D2: loop 5) at the receptor-cytokine interface^23,24^, allowing for high cytokine affinity (type I and type II IL-4Rα:IL-4 *K_D_* = 1 nM; type II IL-4Rα:IL-13 *K_D_* = 20 nM). To investigate the molecular differences between rademikibart and dupilumab, we determined the x-ray crystal structure of rademikibart fragment antigen binding region (Fab) in complex with its target, the IL-4Rα subunit, and compared it to the dupilumab-IL-4Rα binding interface to evaluate how molecular differences in antibody epitopes may impact binding affinity and clinical outcomes.

**Figure 1.**
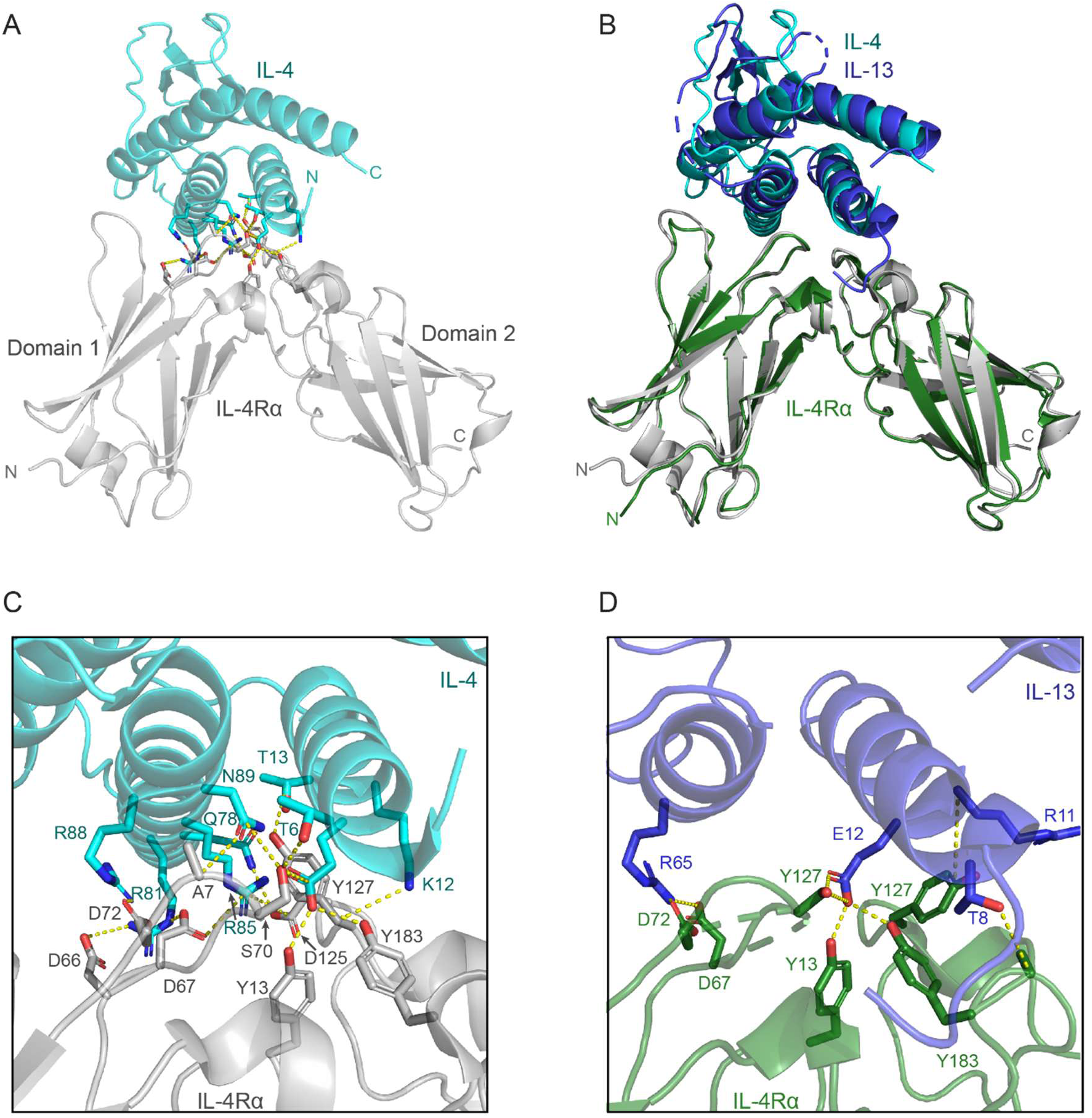
IL-4 and IL-13 bind the IL-4Rα subunit via a conserved mechanism. A. X-ray crystal structure of IL-4 (teal) bound to the IL-4Rα subunit (gray) (PDB ID: 3BP1). Hydrogen bonds and salt bridges between IL-4Rα and IL-4 residues are depicted. B. The IL-13 (dark blue)-IL-4Rα (green) complex (PDB ID: 3BPO) aligned to the IL-4-IL-4Rα complex demonstrates that IL-4 and IL-13 have a conserved binding mode to IL-4Rα that involves both domain 1 (D1) and domain 2 (D2) and the hinge region between D1 and D2. RMSD = 1.132 Å. C. Zoomed in view of the residue-residue interactions between IL-4Rα (grey) and IL-4 (teal). D. Zoomed in view of the residue-residue interactions between IL-4Rα (green) and IL-13 (dark blue).

## Results

To determine the molecular mechanism of rademikibart at atomic resolution, we determined the x-ray crystal structure of the rademikibart Fab in complex with IL-4Rα at a 2.72 Å resolution (**Figure 2A**) with final *R*_work_ value of 23.9% (*R*_free_ 30%). The structure belongs to the C2221 space group with a total of 206 IL-4Rα residues and 429 rademikibart Fab residues (96.8% completeness at the highest resolution shell).

**Figure 2.**
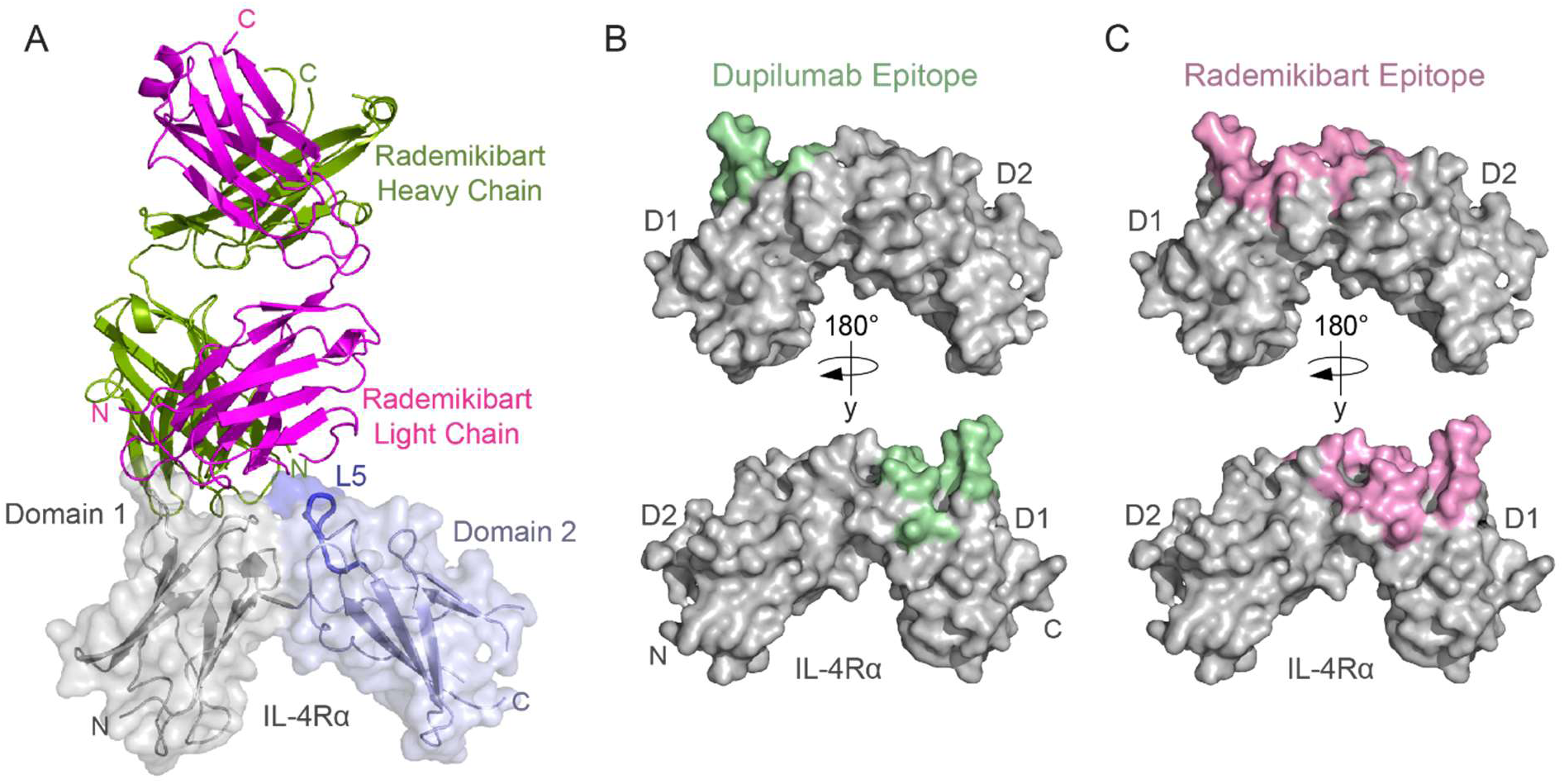
Rademikibart engages the L5 loop of the IL-4Rα subunit. A. Ribbon and overlying surface representation of the x-ray crystal structure of rademikibart bound to the IL-4Rα subunit (2.72 Å resolution). Domain 1 of the IL-4Rα subunit is displayed in grey, domain 2 is displayed in light blue, and the L5 loop is displayed in dark blue. Rademikibart’s heavy (green) and light (magenta) chains are shown in ribbon representation. Note that rademikibart binds across domains 1 and 2 of IL-4Rα. B. Surface representation of the IL-4Rα subunit with the dupilumab epitope (buried surface area 779.8 Å²) displayed in light green. C. Surface representation of the IL-4Rα subunit with the rademikibart epitope (buried surface area 893.3 Å²) displayed in light pink.

To ascertain differences between the rademikibart and dupilumab modes of binding, we compared the rademikibart Fab-IL-4Rα structure to that of the previously determined dupilumab Fab-IL-4Rα complex (PDB ID: 6WGL; 2.82 Å resolution)^25^. The rademikibart and dupilumab epitopes on IL-4Rα were analyzed using PDBePISA^26^. Unlike the dupilumab binding epitope (**Figure 2B**), which only engages loops 2 and 3 of the IL-4Rα D1 domain (residues 66-72 and 90-98) and overlaps the native IL-4 epitope 47% (9/19 residues) and native IL-13 epitope 35% (7/20 residues), the rademikibart epitope (**Figure 2C, Supplemental Table 2**) spans three loops on IL-4Rα D1 and D2, including loop 5 (residues 145-153; **Figure 2A**). Rademikibart’s mode of binding more closely mimics the cognate engagement of the receptor by IL-4 (74% overlap, 14/19 residues) and IL-13 (60% overlap, 12/20 residues, **Supplemental Table 2**). Superposition of the antibody-IL-4Rα co-complexes showed a 54.88° rotation between dupilumab and rademikibart when bound to IL-4Rα (**Figure 3**). This rotation allows for a larger binding interface for rademikibart and facilitates its engagement across the hinge region and to the D2 domain of IL-4Rα, which may potentially increase steric occlusion between rademikibart and IL-4/IL-13 compared to dupilumab.

**Figure 3.**
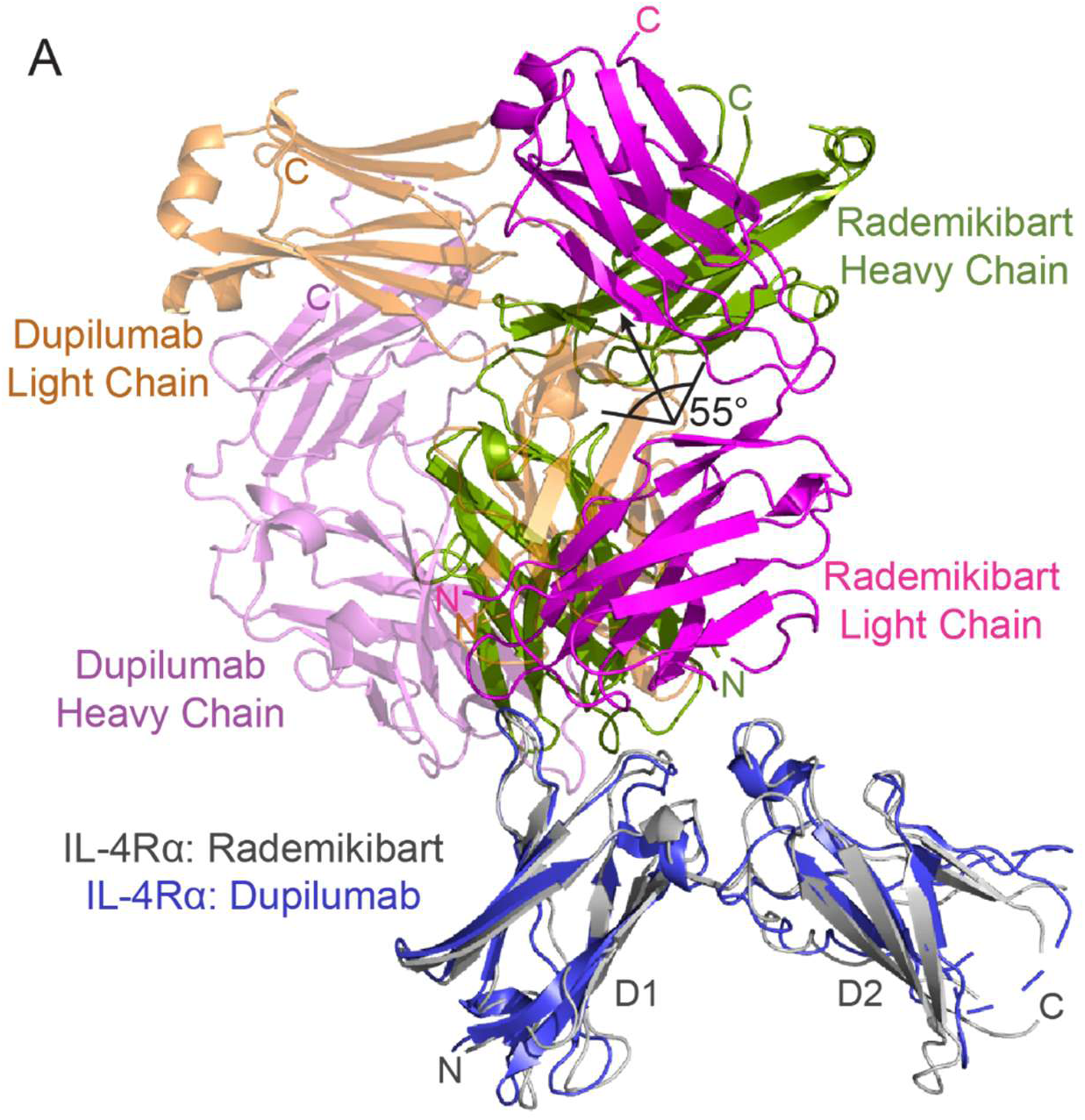
Rademikibart engages a larger surface on IL-4Rα compared to dupilumab. Ribbon representation of the dupilumab Fab-IL-4Rα complex (light chain: orange; heavy chain: purple) with the rademikibart Fab-IL-4Rα complex (light chain: magenta; heavy chain: green) overlaid. Rademikibart displays a 55° rotation about IL-4Rα compared to dupilumab, facilitating engagement across the hinge region to the IL-4Rα L5 loop on D2.

Given the differences in crystallographic antibody binding sites on IL-4Rα and their potential clinical significance, we further investigated and validated the rademikibart Fab-IL-4Rα and dupilumab Fab-IL-4Rα crystal structures using computational co-complex predictions with AlphaFold2^27^. We then performed 400 ns molecular dynamics (MD) simulations of the predicted co-complexes run in triplicate using NAMD for structure relaxation and equilibration (**Supplemental Videos 1-2**)^28^. Overall, there was good agreement between the experimentally and computationally derived antibody-IL-4Rα complexes (rademikibart Fab-IL-4Rα RMSD: 1.3 Å; dupilumab Fab-IL-4Rα RMSD: 2.7 Å; **Figure 4**), confirming their structural mechanisms. Throughout MD simulations, each antibody-receptor complex was found to be structurally stable per low backbone RMSD values (n=3), relative to their mean structures over time (rademikibart Fab-IL-4-Rα: 2.0 ± 0.7 Å; dupilumab Fab-IL-4Rα: 2.5 ± 0.8 Å).

**Figure 4.**
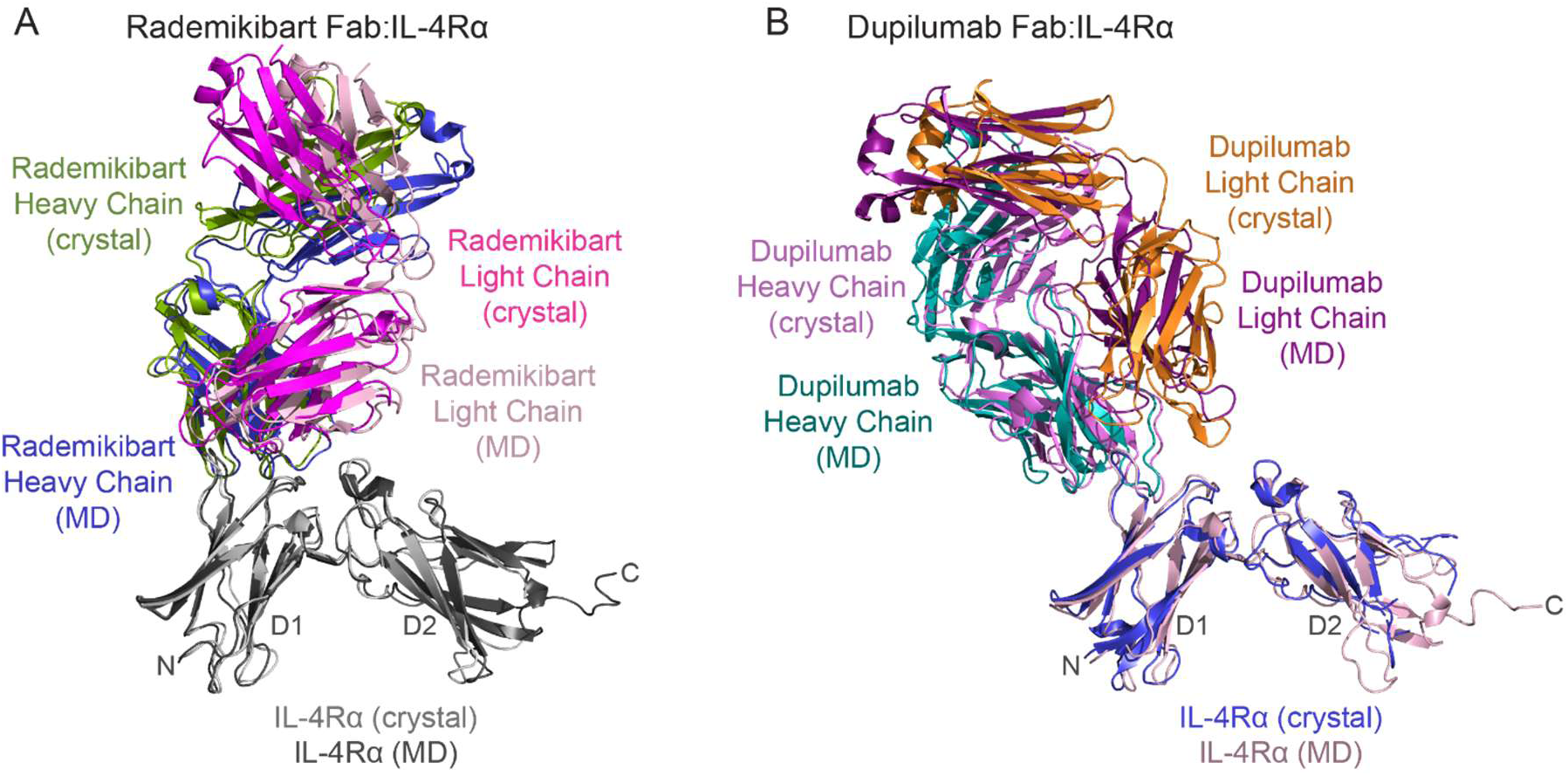
The experimentally and computationally derived rademikibart Fab-IL-4Rα and dupilumab Fab-IL-4Rα co-complexes exhibit good congruence. A. AlphaFold2-derived rademikibart Fab-IL-4Rα co-complex following 400 ns MD simulation overlaid on the x-ray crystallography structure of rademikibart Fab-IL-4Rα from Figure 2A. RMSD = 1.3 Å. B. AlphaFold2-derived dupilumab-IL-4Rα co-complex following 400 ns MD simulation overlaid on the x-ray crystallography structure of dupilumab-IL-4Rα co-complex (PDB ID: 6WGL). RMSD = 2.7 Å.

To further evaluate rademikibart’s interaction with the IL-4Rα D2 domain, B factors, which represent motion or increased protein backbone dynamics, were calculated for each residue (**Figure 5**). Higher B factors indicate a higher degree of motion or dynamics. The L5 loop of the IL-4Rα subunit showed an average B factor of 53.42 Å^2^ for the rademikibart Fab-IL-4Rα complex. In comparison, the average B factor of the L5 loop was 133.22 Å^2^ for the dupilumab Fab-IL-4Rα complex, consistent with a lack of L5 loop engagement in this complex. In addition, the backbone of the L5 loop was positioned further away (>11 Å) from dupilumab than it is from rademikibart (>7.5Å), where notably, contact was maintained between rademikibart and the L5 loop throughout each MD simulation. These data further establish rademikibart’s unique ability to engage the IL-4Rα L5 loop, with an epitope that spans the D1 and D2 domains.

**Figure 5:**
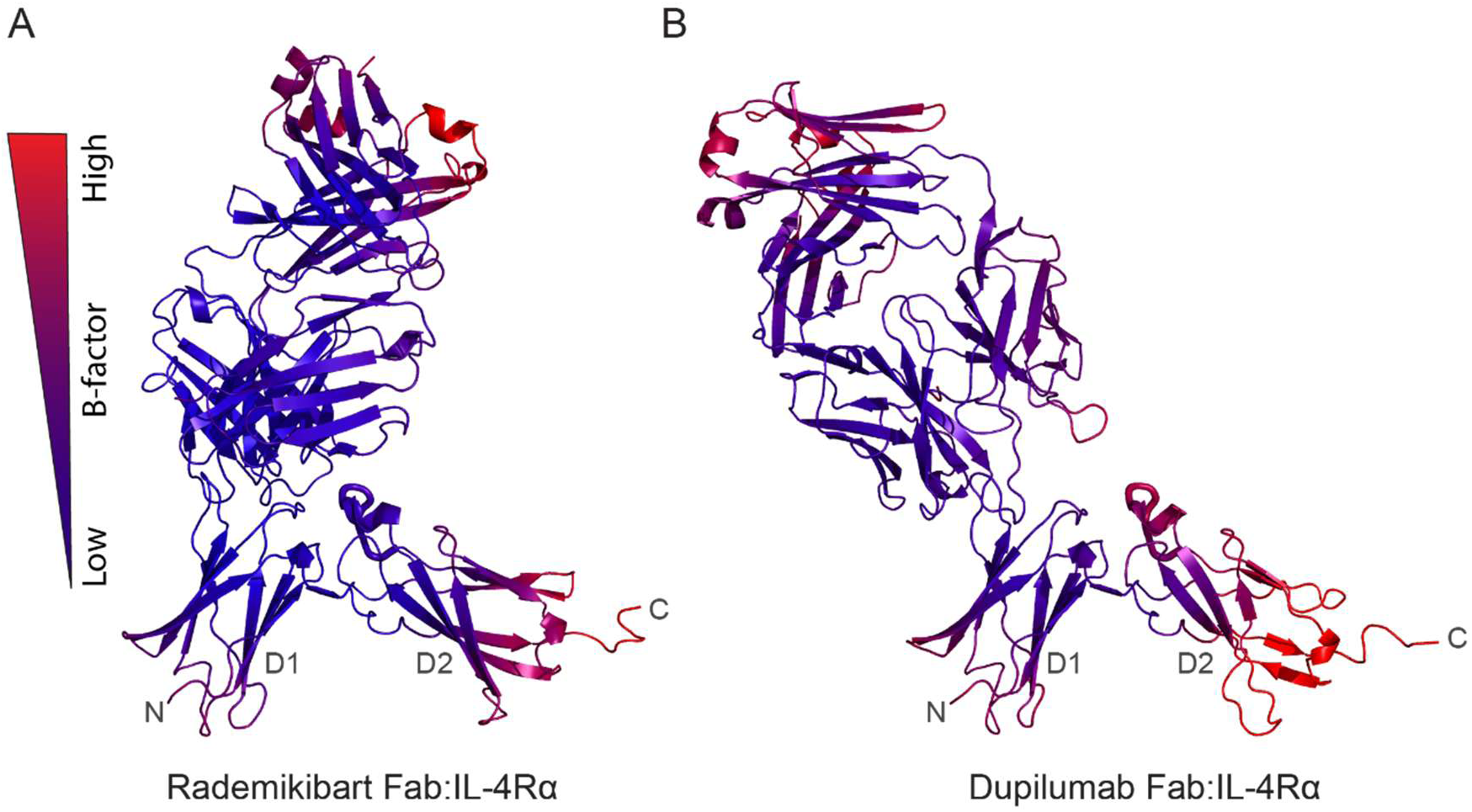
Binding of rademikibart decreases IL-4Rα L5 loop mobility. A. MD equilibrated rademikibart-IL-4Rα co-complex displayed in ribbon format, with B-factors shown in a blue to red gradient, where a more blue color corresponds to less backbone motion (lower B factor) and a more red color corresponds to increased backbone motion (higher B factor). B. MD equilibrated dupilumab-IL-4Rα co-complex displayed in ribbon format, with B-factors shown in a blue to red gradient. Notably, the L5 loop displays higher B-factors in the presence of dupilumab compared to rademikibart, indicating decreased stability of dupilumab-IL-4Rα complex.

Rademikibart has approximately twice the affinity for the IL-4Rα subunit (20.7 pM) compared to dupilumab (45.8 pM)^16^. To better understand these differences, individual inter-residue interactions at the antibody-protein binding interfaces were analyzed for both the x-ray crystal and equilibrated MD co-complex structures. Rademikibart forms several critical intermolecular bonds between itself and the D2 domain, specifically the L5 loop, of IL-4Rα. Y152 of the IL-4Rα L5 loop and Y50 (light chain) of rademikibart participate in π-π stacking interactions, as well as likely hydrogen bonds with rademikibart’s Y50 (light chain) hydroxyl group, seen in both the x-ray crystal and MD structures (**Figure 6A**). In the crystal structure, Y50 (light chain) also participates in π-π stacking interactions with Y101 (heavy chain; **Figure 6A**), whereas in the MD derived structure, Y50 (light chain) forms π-π stacking interactions with IL-4Rα Y152 and H156 (**Figure 6B**). This stabilizing interaction is lost upon a H156D mutation (**Supplemental Figure 1**). IL-4Rα Y152 also forms a hydrogen bond with the R55 (light chain) carbonyl oxygen in the MD-derived structure (**Figure 6C**). Density for R55 is missing in the x-ray crystal structure, however, the flanking rademikibart residues S54 and A56 are positioned similarly to those in the MD-derived structure, suggesting R55 would adopt similar geometry. These interactions would inhibit the cognate hydrogen bonds from forming between Y152 of IL-4Rα and K12 of IL-4 or R11 of IL-13.

**Figure 6:**
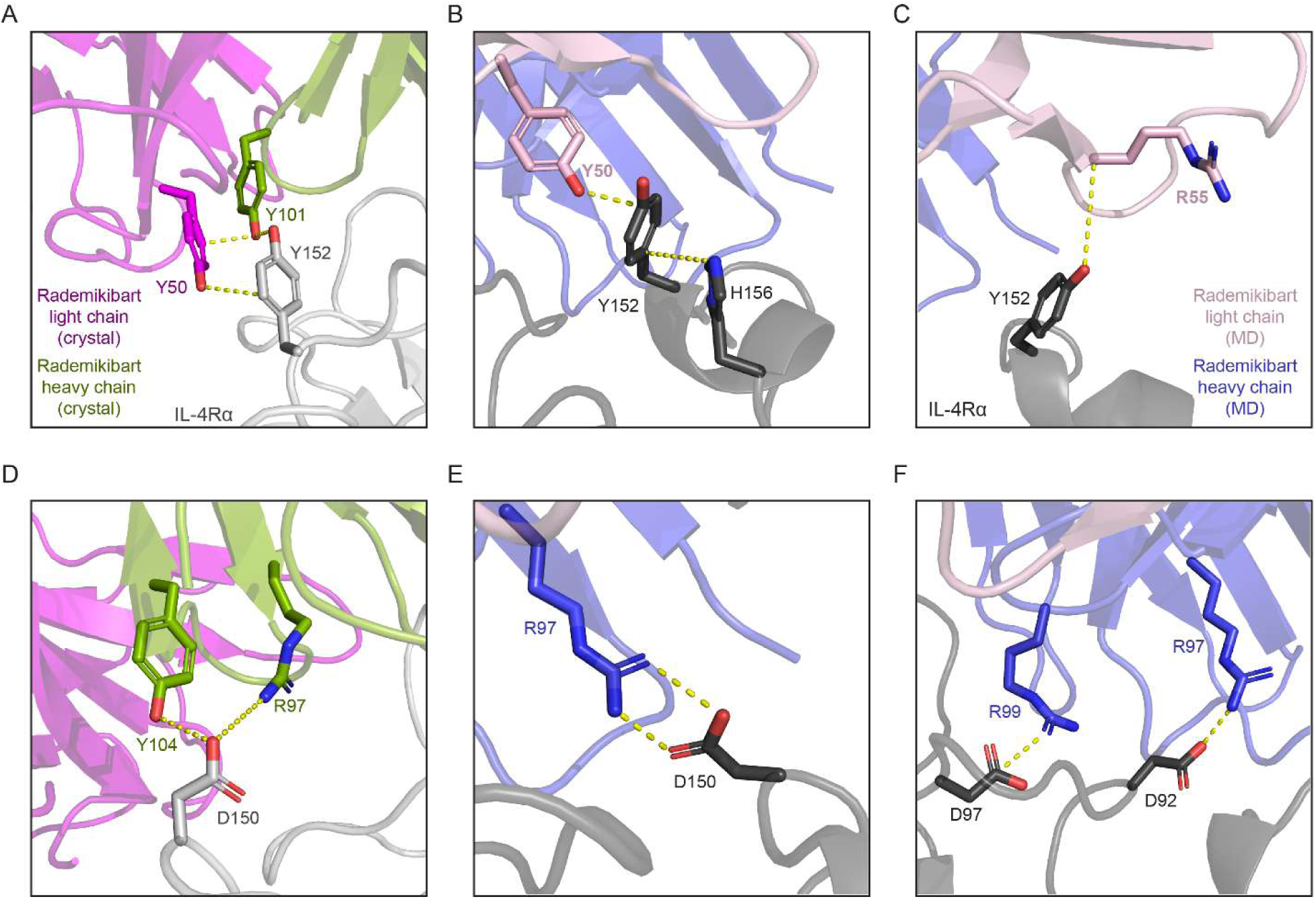
Molecular interactions at the rademikibart-Fab-IL-4Rα interface. A: Y152 of IL-4Rα engages in π-π stacking interactions Y50 (light chain) and Y101 (heavy chain), as well as likely hydrogen bonds between the residues’ hydroxyl groups and side chain carbons per the x-ray crystallography structure. B. Rademikibart Y50 (light chain), IL-4Rα Y152, and IL-4Rα H156 participate in π-π stacking interactions, and likely hydrogen bonds. C. Y152 of IL-4Rα forms a hydrogen bond with the carbonyl oxygen of rademikibart R55 (light chain) per the MD-derived structure. D: IL-4Rα D150 forms a hydrogen bond with rademikibart Y104 (heavy chain), as well as a likely weak or water-mediated hydrogen bond with rademikibart R97 (heavy chain), in the x-ray crystallography structure. E. IL-4Rα D150 forms a strong salt bridge with rademikibart R97 (heavy chain) in the MD-equilibrated structure. F. IL-4Rα D92 and D97 form strong salt bridges and hydrogen bonds with rademikibart heavy chain residues R97 and R99, respectively, per the MD-equilibrated structure.

IL-4Rα’s D150, located within the L5 loop, also hydrogen-bonds with Y104 (heavy chain), with likely formation of a weak hydrogen bond vs. water-mediated hydrogen bond with R97 (heavy chain) (**Figure 6D**). This interaction becomes even more apparent in the equilibrated MD structure, where D150 of IL-4Rα L5 forms strong hydrogen bonds and salt bridges with rademikibart’s R97 (heavy chain; **Figure 6E**). This would prevent D150 of the IL-4Rα from forming its cognate hydrogen bonds with IL-4 Q78 and R85. R97 (heavy chain) and R99 (heavy chain) also form both salt bridges and hydrogen bonds with D92 and D97 of IL-4Rα, respectively (**Figure 6F**). R99 (heavy chain) also forms a hydrogen bond with the carbonyl oxygen of IL-4Rα residue D92.

Rademikibart’s inclusion of Y208 within its epitope would also prevent this residue from being accessible to hydrogen bond with E9 of IL-4 or E12 of IL-13. Conversely, intermolecular interactions between the D2 domain of IL-4Rα with dupilumab are not present. Further, rademikibart engages IL-4Rα residue M39, a residue not found at the dupilumab epitope, via a hydrogen bond with rademikibart R31 (heavy chain; **Supplemental Table 2**). Rademikibart also displays a larger buried surface area (BSA) of 893.3 Å² compared to dupilumab’s 779.8 Å². This is consistent with prior work that has shown a direct correlation between increasing BSA and binding affinity^29^.

## Discussion

We describe the x-ray crystal structure of rademikibart Fab bound to the IL-4Rα subunit, providing mechanistic detail on how this next-generation biologic engages IL-4Rα uniquely and more optimally than a first-generation IL-4Rα inhibitor. Strikingly, rademikibart engaged both the D1 and D2 domains of IL-4Rα, enabling rademikibart’s epitope to closely overlap with the natural binding sites of IL-4 and IL-13 on IL-4Rα, which contrasts with dupilumab. Similar to dupilumab, we identified that rademikibart binds the L2 and L3 loops of the IL-4Rα D1 domain; however, rademikibart also binds the L5 loop of the IL-4Rα D2 domain, which dupilumab does not. D150 and Y152 within the IL-4Rα L5 loop are not highly conserved residues when compared to proteins with homologous sequences from a variety of species per Consurf analysis^30–35^ (**Supplemental Table 3**). This lack of conservation introduces significant divergence in the human rademikibart epitope location and likely contributes to the antibody’s selectivity for only the human IL-4Rα^16^. Given IL-4 and IL-13 engage both the D1 and D2 domains of IL-4Rα, we expect that occlusion of both interfaces provides superior inhibition of IL-4/IL-13 signaling (**Supplemental Video 3:** https://vimeo.com/1082000394).

Dupilumab has incomplete engagement of the full native IL-4/IL-13 epitope on IL-4Rα. Conversely, rademikibart’s 54.88° rotation from dupilumab when in complex with IL-4Rα allows for binding to both IL-4Rα domains. This provides a more stable interaction with IL-4Rα as evidenced by rademikibart’s superior binding affinity and improved co-complex stability as demonstrated by MD simulations. This correlates with in vitro biochemical studies showing that rademikibart displays greater inhibition of downstream STAT6 activation^16^. Together, these results provide a molecular rationale as to why rademikibart is an optimized, next-generation IL-4Rα-blocking antibody with higher-affinity epitope engagement and potent inhibition of IL-4– and IL-13–mediated signaling pathways compared with dupilumab. The structural differences between rademikibart and dupilumab also provide a mechanistic rationale for differences observed both preclinically and in the clinic.

Across preclinical mechanistic studies, rademikibart consistently induced strong suppression of canonical STAT6 phosphorylation, TF-1 proliferation, and TARC (CCL17) production, frequently surpassing the potency benchmarked with dupilumab^16^. At the structural-immunobiology level, IL-4Rα exists in a dynamic equilibrium between surface-resident receptor and endosomal pools, undergoing both constitutive endocytosis and ligand-accelerated internalization—processes that tightly regulate the amplitude and duration of JAK/STAT signaling outputs^36,37^. Comparative internal data indicate that rademikibart engages an epitope positioned closer to the IL-4Rα D1–D2 hinge, a region that governs conformational flexibility required for heterodimer assembly, whereas dupilumab binds a more lateral D1 epitope. This difference in binding geometry translates into markedly different receptor trafficking behaviors: rademikibart accelerates IL-4Rα internalization approximately 2-fold over baseline and ∼1.7-fold more than dupilumab (data on file, Connect Biopharma), resulting in greater and more rapid receptor down-modulation. Functionally, this enhanced internalization shifts the signaling landscape by suppressing sustained STAT6 activation more efficiently and by altering survival-related signaling nodes. From a safety perspective, this may play an important role in differences observed between rademikibart and dupilumab in the clinic.

Dupilumab has repeatedly been linked to eosinophil–associated disorders—including hypereosinophilia, eosinophilic pneumonia, and EGPA^11,13,14^—as well as facial erythema/head–and–neck dermatitis^10,12^ and ocular surface disease such as conjunctivitis and keratitis^6–9^. Rademikibart has produced large improvements in T2-predominant disease activity without triggering treatment-emergent eosinophilia or hypereosinophilia (>3000 cells/µL)^38–40^. In asthma patients, mean eosinophil levels were reduced following rademikibart treatment, and the incidence of post-baseline elevations >1500 cells/µL was comparable to or lower than placebo^41^.

Receptor-level biology supports a mechanism by which rademikibart avoids treatment-emergent eosinophilia and eosinophil–associated disorders. Internal mechanistic assays show that rademikibart induces a 3.5–4-fold increase in eosinophil apoptosis relative to IL-4 baseline–induced apoptosis, whereas dupilumab produces only a ∼1.3–1.5-fold increase under matched assay conditions (data on file, Connect Biopharma). These quantitative differences in receptor turnover and survival signaling provide a structural-mechanistic explanation for how rademikibart counterbalances IL-4/IL-13–driven trafficking forces that would otherwise transiently increase circulating eosinophils. Mechanistically, this is consistent with evidence that sufficiently high IL-4 exposure can directly induce eosinophil apoptosis^42^, and that the rate and efficiency of IL-4Rα internalization critically govern temporal signaling outputs^36,37^. Integrating these observations, rademikibart’s unique receptor-modulating properties provide a coherent mechanistic rationale for the absence of treatment-emergent eosinophilia—differentiating it from dupilumab where receptor trafficking or signaling perturbations may transiently elevate circulating eosinophils. Clinically, rademikibart demonstrates meaningful and durable efficacy across type 2 (T2) inflammatory diseases, with signals that compare favorably to historical dupilumab benchmarks while reflecting distinct receptor-level pharmacology.

In Atopic Dermatitis (AD), the SEASIDE CHINA randomized phase II trial with rademikibart monotherapy produced rapid, clinically meaningful improvements in eczematous lesions, pruritus, and quality of life during the initial 16-week treatment period, with continued deepening of response through the 52-week study duration^39^. Among patients who achieved EASI-50 at week 16 and entered Stage 2, responses continued to strengthen through week 52, with similar efficacy observed in both biweekly (Q2W) and monthly (Q4W) maintenance dosing with EASI-75 responses reaching 84.6% and 84.8%, and ≥4 point PP-NRS reductions were achieved in 60.4% and 70.3% of patients in the respective groups. Among those who had already improved by week 16, 90.1% (Q2W) and 92.3% (Q4W) and 85.3% and 94.8% maintained their EASI-75 response and ≥4 point PP-NRS reduction respectively^39^. Similar findings from the phase III RADIANT-AD trial further confirm this enhanced efficacy. In a recent late-breaking oral presentation at the American Academy of Dermatology 2026 annual meeting, rademikibart demonstrated significantly greater clinical responses versus placebo at Week 16, with 74.2% of treated participants achieving EASI-75 compared with 34.4% in the placebo group, and 47.7% achieving IGA 0/1 versus 17.6%, respectively. Clinically meaningful itch improvement was also observed, with a ≥3-point reduction in peak pruritus NRS achieved by 54.7% of rademikibart-treated patients versus 27.5% on placebo at Week 16. Responses continued to deepen over time, such that by Week 52, EASI-75 and IGA 0/1 response rates reached approximately 96% and 87%, respectively, and itch responses exceeded 90% among patients receiving continuous rademikibart treatment or those switched from placebo to rademikibart 300 mg every 2 weeks after Week 16^40^.

A mechanistic rationale for the deep skin clearance and sustained durability of response observed with rademikibart can be inferred from rademikibart’s structural properties, as its D1-D2 spanning epitope and high affinity IL-4Rα engagement promotes more effective receptor occupancy and significantly enhanced IL-4Rα internalization, resulting in a greater and more sustained suppression of IL-4/IL-13 signaling. This pharmacologic durability provides a mechanistic explanation for the clinical finding that Q4W maintenance dosing preserved—or even improved—Week 16 responses at levels comparable to Q2W dosing in SEASIDE CHINA^39^. In contrast, dupilumab responses in phase II and III trials typically plateau by week 16, with diminishing incremental benefit during extended treatment, and cannot support Q4W dosing without loss of efficacy^15,43^. Together, the structural advantages of rademikibart and the clinical trajectory observed in SEASIDE CHINA and RADIANT-AD indicate that rademikibart provides strong, durable disease control with dosing flexibility that may offer meaningful advantages over existing IL-4Rα–targeted biologics for patients with moderate to severe AD.

A global phase 2b trial of moderate-to-severe asthma with T2 inflammation showed rademikibart produced significant improvements in lung function and asthma exacerbations, with early signs of clinical improvement, detectable as early as week 1, via objective measures including evening peak expiratory flow (PEF) and clinically meaningful gains in forced expiratory volume (FEV₁) over the treatment period^38^. Notably, an exploratory post hoc analysis of the phase 2 trial demonstrated that 77% of the week-1 FEV₁ improvement was achieved within 24 hours after the first subcutaneous 600-mg loading dose of rademikibart, corresponding to a +93 mL pre-bronchodilator (BD) FEV₁ change from baseline vs –101 mL for placebo^44^. These findings highlight a rapid onset profile not observed in dupilumab asthma programs, where FEV₁ gains typically emerge over subsequent weeks. Mechanistically, this accelerated clinical response aligns with rademikibart’s structural advantages: its hinge-proximal IL-4Rα epitope enforces a more restrictive receptor conformation, drives approximately 2-fold higher IL-4Rα internalization rates, and produces more significant STAT6 suppression than dupilumab. Together, these receptor-level dynamics create a more immediate attenuation of IL-4/IL-13 signaling, providing a structural-biological basis for the near-immediate (within 24 hours) improvements in airflow and lung function observed clinically^44^.

The rapid onset of clinical efficacy observed in the aforementioned study has informed the design of ongoing acute-care trials including: a phase 2 trial in acute asthma exacerbations (SEABREEZE STAT Asthma, NCT06940141) evaluating add-on rademikibart (single-dose adjunct to standard care) and a phase 2 trial in acute COPD exacerbations (SEABREEZE STAT COPD, NCT06940154) targeting T2-eosinophilic phenotypes in urgent-care settings. These programs operationalize the hypothesis that faster, deeper suppression of IL-4/IL-13 signaling can translate into short-term physiologic benefits for patients during exacerbations. This hypothesis is further supported by a recent company-reported intravenous (IV) study of rademikibart 300 mg demonstrating early, clinically meaningful improvements in lung function (FEV₁) in patients with asthma or COPD, observed as early as 15 minutes following IV dosing, consistent with rapid IL-4Rα pathway engagement^45^ .

As we continue to develop new and more precisely targeted therapeutics, it is important to understand where and how these drugs specifically engage their targets to advance our mechanistic insight, enable structure-based drug optimization, and inform prescription choices. Our molecular data provide a structural rationale for rademikibart’s enhanced IL-4Rα inhibition—driven by high-affinity epitope engagement of the D1–D2 hinge and superior IL-4/IL-13 receptor internalization kinetics—consistent with the clinical pattern of faster and deeper responses in AD and asthma, further motivating evaluation of rademikibart use for other T2 inflammatory conditions, including acute asthma and COPD exacerbations.

## Methods

### Crystallization, data collection, and structure determination of the rademikibart Fab-IL-4Rα complex

Crystals were grown using sitting drop vapor diffusion at 20 °C. A volume of 200 nL of 10.7 mg/mL rademikibart Fab-IL-4Rα complex was mixed with 200 nL of reservoir solution containing 0.1M SPG buffer (pH 5) and 25% w/v PEG 1500. Crystals were flash-frozen using liquid nitrogen after transfer to a cryo-protective solution (0.1M SPG buffer (pH 5), 25% w/v PEG 1500, and 25% Glycerol).

Multiple data sets were then collected at beamline BL17U1 at SSRF using an Eiger 16M detector. A total of 260° of data were collected in 0.5° increments with a detector distance of 250 mm. All data were then processed using HKL2000. Molecular replacement using Phaser was completed with the model of IL-4Rα from the structure contained in Protein Data Bank Code 3BPL. Structure refinement was performed using Refmac5. Representative images of the rademikibart Fab-IL-4Rα complex were then generated using PyMOL (3.1.3.1).

### Co-complex structure predictions and molecular dynamics simulations

The structure of the rademikibart-IL-4Rα binding complex was predicted using AlphaFold2^27^. The crystal structure of the dupilumab-IL-4Rα with resolution of 2.82 Å was obtained from the PDB website (ID: 6wgl)^25^. Schrödinger Maestro was used for the protein preparation work: capping terminals, building disulfide bonds, determining protonation states with PROPKA and minimizing the structure with restraint to 0.3 RMSD in the OPLS force field. The structures were placed in water boxes with a 15 Å cushion around the complex, and Na+ and Cl- ions were added at a concentration of 0.15 M to maintain the neutrality of the systems. The tLEaP program from the AmberTools23 package^46–48^ was used to generate the parameter and topology input files for the MD simulations. MD simulations were then run in triplicate using NAMD. The system was equilibrated at 310 K in three steps before the production run: (i) minimization of the solvent, (ii) minimization of the solvent and protein side chains, and (iii) minimization of the entire system. For the production run, Hydrogen Mass Repartitioning (HMR) was used, allowing the MD simulations to be conducted with a 4 fs time step, with a total simulation duration of 400 ns. X-ray maps were calculated from the atom coordinates obtained from MD trajectories using the sfall program from the CCP4 package^49^, and the structures were fitted to the map using real-space refinement in Phenix^50^.

### Epitope identification and analysis

The x-ray crystallographic or final molecular dynamics simulation structures were used as input into PDBePISA, where epitopes for IL-4, IL-13, rademikibart, and dupilumab were identified^26^. Residues involved in intermolecular interactions were also identified using PDBePISA, as well as visual inspection via PyMOL (version 3.1.3.1)^51^. The buried surface area at each interface was determined used PDBePISA. Hydrogen bond occupancies at the antibody-protein interfaces were calculated using the Hydrogen Bonds plugin in VMD^52^. The percent overlap of the rademikibart and dupilumab epitopes with those of the IL-4/IL-13 binding sites were then calculated.

### Data analysis and visualization

Data visualization was performed using PyMOL (version 3.1.3.1)^51^ and final figures were generated using Adobe Illustrator.

## Author Contributions Statement

Conceptualization: CB, RC; Data curation: YS, KN, MH, RC, CGB; Formal analysis: YS, KN, MH, CGB; Funding acquisition: CGB; Investigation: YS, KN, MH, RC, CGB; Project administration: RC, CGB; Supervision: RC, VSB, CGB; Validation: YS, KN, MH, VSB, RC, CGB; Visualization: YS, KN, MH, CGB; Writing-original draft: YS, KN, MH; Writing-review and editing: YS, KN, MH, VSB, RC, CGB. CGB is the guarantor of the work.

