## supplemental_tables_figure for "Crystal structure and molecular dynamics simulations of rademikibart Fab-IL-4Rα complex reveal biochemical basis for next-generation potent IL-4Rα inhibition in type 2 allergic and inflammatory diseases"

Yuanjun Shi^1*^, Kelsey Nolden^2*^, Minh Ho^3^, Haote Li^1^, Victor S. Batista^1^, Raúl Collazo^4^, Christopher G. Bunick^3,5,6^

| Rademikibart Fab-IL-4Rα Co-complex | |
| --- | --- |
| Resolution range | 44.21 - 2.722 (2.819 - 2.722) |
| Space group | C 2 2 21 |
| Unit cell | 96.001 164.5 238.109 90 90 90 |
| *Data processing* | |
| Total reflections | 8540935 |
| Unique reflections | 50589 (4859) |
| Multiplicity | 7.4 (7.2) |
| Completeness (%) | 99.44 (96.83) |
| Mean I/sigma(I) | 9.211 (1.066) |
| Wilson B-factor | 64.49 |
| R_meas_ | 0.169 (2.349) |
| R_pim_ | 0.085 (1.202) |
| CC_1/2_ | 0.988 (0.37) |
| *Refinement* | |
| Reflections used in refinement | 50577 (4859) |
| Reflections used for R_free_ | 2648 (242) |
| R_work_ | 0.2394 (0.3986) |
| R_free_ | 0.3004 (0.4133) |
| Number of non-hydrogen atoms | 9601 |
| Macromolecules/ligands/solvent | 9450/53/98 |
| Protein residues | 1227 |
| RMS(bonds)/(angles) | 0.012/1.63 |
| Average B-factor | 95.71 |
| Macromolecules/ligands/solvent | 95.92/133.23/55.17 |
| *Ramachandran, rotamer, and clash statistics* | |
| Ramachandran favored (%) | 91.45 |
| Ramachandran allowed (%) | 7.88 |
| Ramachandran outliers (%) | 0.66 |
| Rotamer outliers (%) | 11.39 |
| Clashscore | 9.57 |
| Average B-factor | 95.71 |
| macromolecules | 95.92 |
| ligands | 133.23 |
| solvent | 55.17 |

**Supplemental Table 1**. X-ray crystal parameters table. Statistics for the highest-resolution shell are shown in parentheses.

| **IL-4** | **IL-13** | **Dupilumab** | **Rademikibart** |
| --- | --- | --- | --- |
| Y38 | Y38 | - | - |
| - | - | - | M39 |
| - | - | - | S40 |
| - | - | Y62 | - |
| - | - | Q63 | - |
| L64 | L64 | L64 | L64 |
| - | - | - | V65 |
| F66 | F66 | F66 | F66 |
| L67 | - | L67 | L67 |
| - | - | L68 | L68 |
| - | - | S69 | S69 |
| - | - | E70 | E70 |
| - | - | A71 | - |
| - | - | H72 | H72 |
| - | - | L89 | - |
| - | - | M90 | M90 |
| D91 | - | D91 | D91 |
| D92 | D92 | D92 | D92 |
| V93 | V93 | - | - |
| V94 | V94 | V94 | V94 |
| S95 | S95 | - | S95 |
| A96 | A96 | - | A96 |
| D97 | D97 | D97 | D97 |
| - | - | - | N98 |
| - | - | Y99 | - |
| S118 | S118 | - | - |
| - | E119 | - | - |
| - | H120 | - | - |
| - | - | - | P149 |
| D150 | D150 | - | D150 |
| N151 | - | - | - |
| Y152 | Y152 | - | Y152 |
| N155 | N155 | - | N155 |
| H156 | H156 | - | - |
| - | Q206 | - | - |
| C207 | C207 | - | - |
| Y208 | Y208 | - | Y208 |
| - | N209 | - | - |

**Supplemental Table 2.** Residues belonging to the epitopes of IL-4, IL-13, dupilumab, and rademikibart on IL-4Rα. The dupilumab epitope overlaps the cognate IL-4/IL-13 epitopes by 30.4% (7/23 residues) whereas the rademikibart epitope overlaps by 56.5% (13/23 residues).

| Res # | Seq | Score |
| --- | --- | --- |
| 145 | N | -0.616 |
| 146 | P | -0.543 |
| 147 | Y | -1.337 |
| 148 | P | -0.392 |
| 149 | P | 2.459 |
| 150 | D | 2.473 |
| 151 | N | 0.524 |
| 152 | Y | 2.478 |
| 153 | L | -1.173 |

**Supplemental Table 3.** Conservation scores of the IL-4Rα L5 loop (residues 145-153) as determined by Consurf analysis. A more negative number indicates a higher degree of conservation compared to sequence-derived homologues of different species, whereas a more positive number indicates a lower degree of evolutionary conservation.

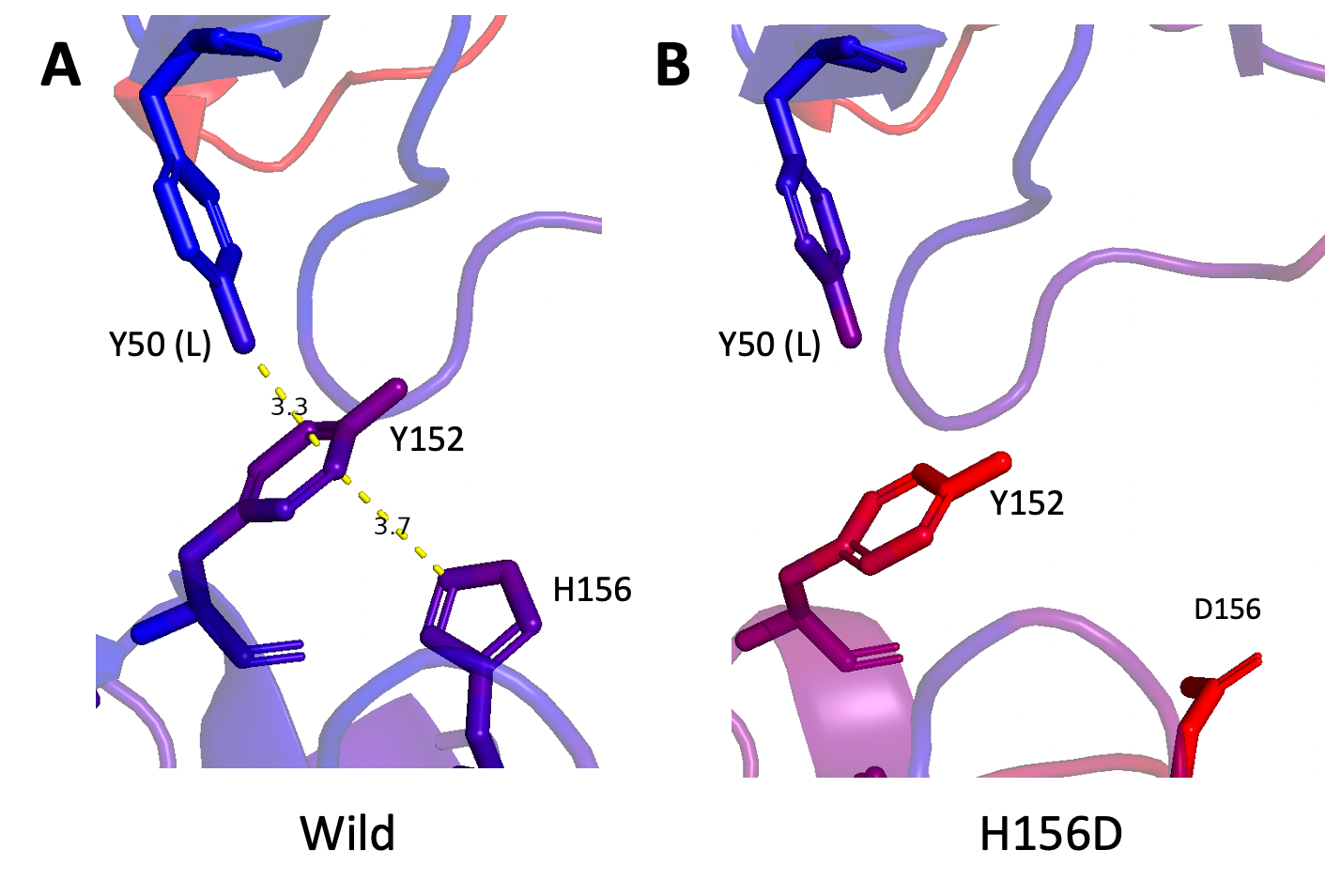

**Supplemental Figure 1. A H156D mutation of IL-4Rα disrupts a critical intermolecular interaction.** A. Wild-type IL-4Rα bound to rademikibart Fab exhibits π-π stacking interactions between rademikibart Y50 (light chain), IL-4Rα Y152, and IL-4Rα H156 per the MD-equilibrated structure. B. A H156D mutation of IL-4Rα causes disruption of the native in π-π stacking interaction, decreasing rademikibart Fab-IL-4Rα complex stability.
